# Dispersed Sleep Microstates and Associated Structural Changes in GBA1 Mouse: Relevance to Rapid Eye Movement Behavior Disorder

**DOI:** 10.1101/2021.05.26.445845

**Authors:** Cigdem Gelegen, Diana Cash, Katarina Ilic, Millie Sander, Eugene Kim, Camilla Simmons, Michel Bernanos, Joana Lama, Karen Randall, Jonathan T. Brown, Svjetlana Kalanj-Bognar, Samuel Cooke, K Ray Chaudhuri, Clive Ballard, Paul Francis, Ivana Rosenzweig

**Affiliations:** Sleep and Brain Plasticity Centre, Department of Neuroimaging, Institute of Psychiatry, Psychology and Neuroscience (IoPPN), King’s College London (KCL), UK; Basic and Clinical Neuroscience, IoPPN, KCL, UK; BRAIN, Department of Neuroimaging, KCL, UK; University of Zagreb School of Medicine, Zagreb, Croatia; College of Medicine and Health, University of Exeter, Exeter, UK; Institute of Psychiatry, Psychology & Neuroscience, Wolfson Centre for Age-Related Diseases, Guy’s Campus, King’s College London, UK; King’s College London and Parkinson’s Foundation Centre of Excellence, King’s College Hospital, London, London, United Kingdom; Sleep Disorders Centre, GSTT, London, UK

**Keywords:** GBA1, neuroimaging, sleep, mouse model, RBD, LBD

## Abstract

Rapid eye movement (REM) sleep behaviour disorder (RBD) is a rare parasomnia that may predict the later occurrence of alpha-synucleinopathies. Variants in the gene encoding for the lysosomal enzyme glucocerebrosidase, GBA, strongly increase the risk of RBD. In a GBA1-mouse model recently shown to mimic prodromal stages of α-synucleinopathy, we now demonstrate striking REM and NREM sleep abnormalities accompanied by distinct structural changes in the more widespread sleep neurocircuitry.

---

Idiopathic rapid eye movement (REM) sleep behaviour disorder (iRBD) may present the prodromal stage of an α-synucleinopathy^1^. Variants in the gene encoding for the lysosomal enzyme glucocerebrosidase, GBA, strongly increase the risk of iRBD*^2^*. The rate of conversion to neurodegeneration is also increased, and may be faster among severe GBA variant carriers.

To date, surprisingly little is still known about the macroscopic and microscopic sleep architecture in iRBD^3, 4, 5, 6^. Even less is known about its relationship with the GBA-genotype and any associated neuroanatomical or circuitry changes^7^. In the past, it has been argued that a breakdown in the specfic “REM-on” brainstem sleep circuitry residing in the region of the sublaterodorsal tegmental nucleus and the coeruleus/subcoeruleus locus (SLD)^8^, i.e. the structure analogous to the peri-locus coeruleus-alpha in cat and sublaterodorsalis nucleus (e.g. A6sc) in rodents, might lead to RBD-associated sleep architectural changes^3, 9, 10, 11^. However, the specific changes have by and large remained elusive^11^. In order to put this concept to the test, whilst accounting for the effect of GBA1-genotype, we used a GBA1-mouse model (heterozygous D409V/WT), recently shown to mimic early prodromal stage of α-synucleinopathy^12^.

*Firstly, in order to explore putative similarities of macroscopic and microscopic sleep structure in* D409V/WT mice^12^ *to those previously reported in iRBD*^3, 4, 5^, 24-hour video-EEG recordings of aged-matched D409V/WT and wild type (WT) mice were undertaken. Increased (sleep) latency to NREM, longer and deeper bouts of NREM, overall reduced amount of REM sleep, accompanied by a distinct change in REM’s theta power and range, were all found in D409V/WT by comparison to WT mice (Figure 1a–e).

**Figure 1.**
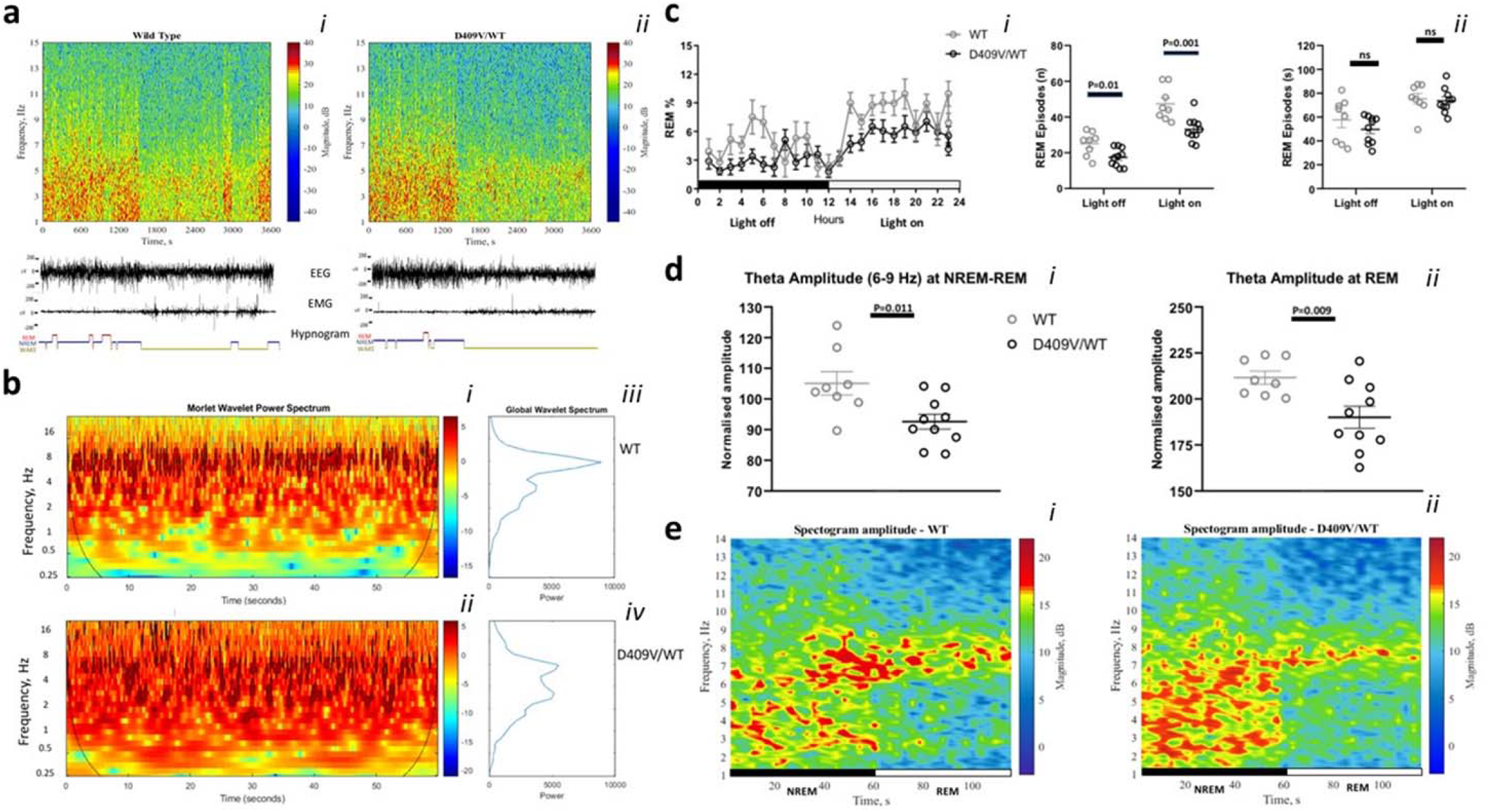
Specificity of Rapid Eye Movement (REM) sleep in D409V/WT. In <u>**a**</u>, **the representative EEG power, EEG/EMG traces, and hypnograms over one-hour period are shown**. Examples of amplitude spectograms (*right*, scale-bar;dB), EEG/EMG traces, and hypnograms over one-hour period at “light-on” are shown from a representative WT and D409V/WT-mouse. <u>In **b**</u>, **Morlet-and-Global-Wavelet-Spectrums of REM sleep in WT and D409V/WT-mice are depicted**. Morlet spectrums (*right*, scale-bar;log10 relative power) were generated by averaging all REM-periods lasting one minute at “lights-on”-period; shown as a function of frequency and time(b*i,bii*). A significantly reduced power at the thetafrequency-range and an intrusion of a lower-frequency-range(1-4 Hz,delta) in D409V/WT is shown(b*i,bii*). Global-wavelet-spectrums similarly depict changes(b*iii,biv*). <u>In **c**</u>, a significant reduction in REM in D409V/WT, by comparison to WT, during 24-hours-EEG-recordings are shown(c*i*;*P*=.035). The decrease was due to a reduction in the number(*P*=.001 and *P*=.01;c*ii*)and not duration of REM-episodes(c*iii*). <u>In **d**</u>, **significant reductions in theta range are shown at both NREM-to-REM-transitions**(60s;*P*=.011;d*i*) **and during the subsequent REM**(60-120s;*P*=.009;d*ii*). I<u>n **e**</u>, **similar changes by comparison to WT are depicted via representative spectograms of EEG-data-power**(dB); shown as a function of frequency and time(e*i*,e *ii*).

Specifically, the time spent in NREM throughout the 24-hour recording was similar between the two groups (Supplement **Figure S3**). D409V/WT mice displayed a longer latency to NREM during the twelve hours lights-on (the “sleep” period) (361.6 ± 24.76 vs 505. ± 52.19 s; *P* = .035, t = 2.29, DF = 16; *t*-test) (**Figure S3b**). However, once in NREM, D409V/WT mice remained in this state longer, as shown by longer duration of the average NREM episodes during the “sleep” period (296.5 ± 20.77 vs 357.2 ± 16.35 s; *P* = .033, t = 2.33, DF = 16; *t*-test) (**Figure S3c**). Accordingly, sleep fragmentation and the overall number of arousals were similarly reduced (172.8 ± 10.67 vs 137.4 ± 3.53; *P* = 0.0024, t = 3.56, DF = 17; *t*-test) (**Figure S3d,e**)

Strikingly, D409V/WT mice also spent significantly less time in REM (*P*(genotype)_RM-**two**-**wayANOVA**_ = .035; F(genotype)_1,9_ = 6.098; Figure 1c), predominantly due to a reduction in the number of REM episodes (lights-on: 47.38 ± 3.32 vs 33.10 ± 2.19; t = 3.71, *P* = .001; lights off: 25.13 ± 2.34 vs 17.40 ± 1.61, *P* = .0012, t = 2.79; DF = 16; *t*-test), rather than their duration (Figure 1c).

Further spectral EEG analyses indicated a significant reduction in the theta frequency (6-9 Hz) both at the NREM-REM transitions (105.1 ± 3.76 vs 92.61 ± 2.45, *P* = .011, t = 2.87, DF = 16; *t*-test) and during the subsequent REM (211.6 ± 3.52 vs 190.1 ± 5.91, *P* = .009, t = 2.92; DF = 16; *t*-test) (Figure 1b,d,e).

*Secondly, we set to investigate how GBA genotype might affect the (sleep) neurocircuitry* by conducting a high resolution *ex-vivo* magnetic resonance imaging (MRI) of D409V/WT, homozygous (D409V/D409V) and WT mice (Figure 2;**S4-7**).

**Figure 2.**
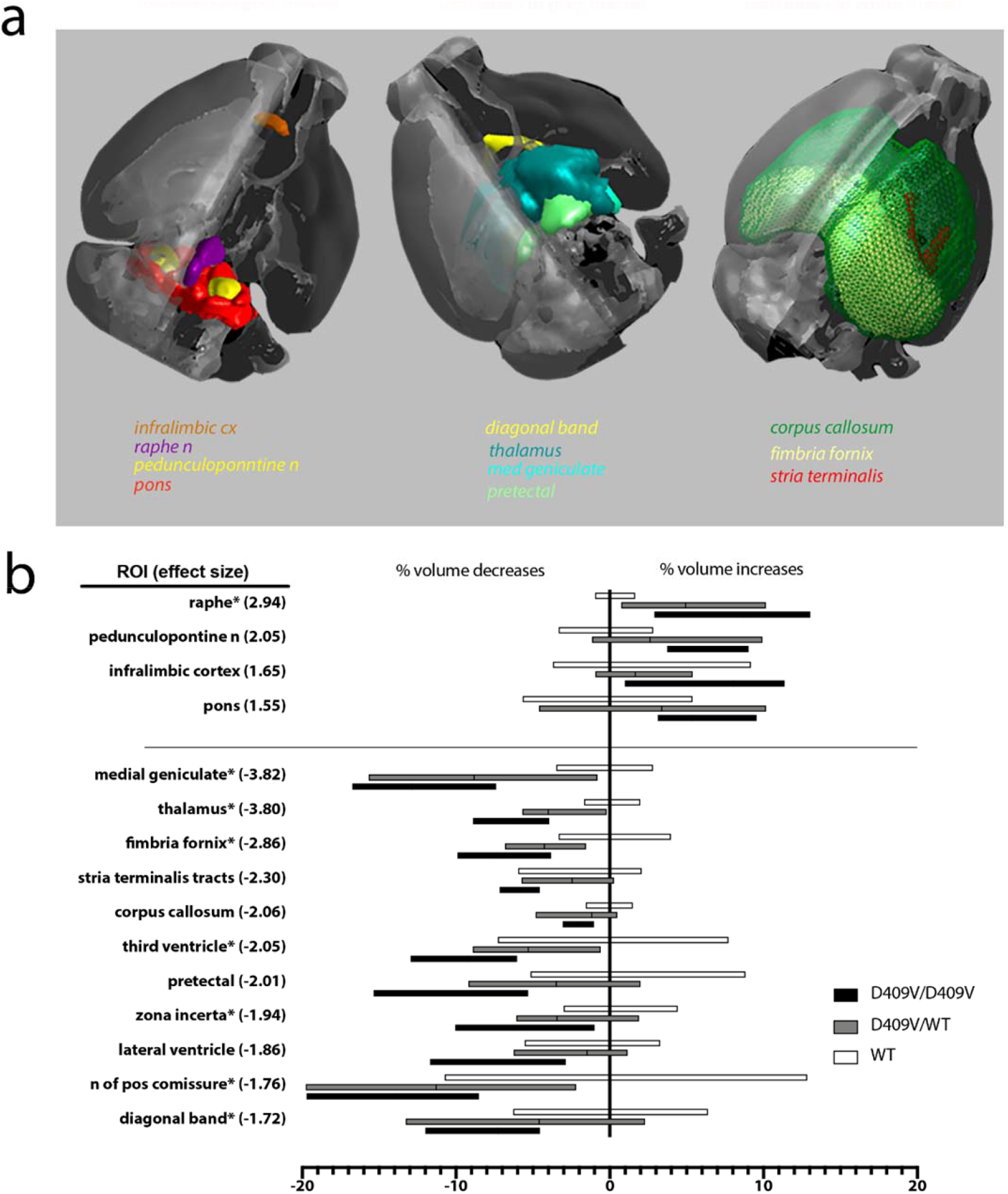
GBA1 Genotype-Driven Neuroanatomical Phenotypes. ROIs (a, b) with significant differences (*P*<.05, FDR corrected) in volume, expressed as % of the averaged wildtype group. Note relative volumes decrease with genotype (WT>D409/WT>D409/D409), except in the dorsal raphe nucleus, pedunculopontine nuclei and infralimbic cortices where there is an opposite trend (WT<D409/WT<D409/D409).

Structural comparisons have highlighted GBA-genotype-driven hypotrophic changes in widespread cortical, subcortical and white matter regions (Figure 2). Comparative enlargements were, however, also noted. For example, hypertrophic plastic changes were conspicuous in the infralimbic cortices, the pons, pedunculopontine (PPT) and the dorsal raphe nuclei (DRN; Figure 2; **S4-7**). Of note, preliminary immunochemistry findings suggest that hypertrophic changes in the PPT, albeit not in DRN, could be, at least in part, driven by the glial proliferation (Supplement **FigureS8a**; *Iba1* (mean±S.D.): WT: 138.6 ± 8.29; D409V/WT: 168.3 ± 17.78: D409V/D409V: 181 ± 36.050; *P*_D409V/WTvsWT_ = .014 and *P*_D409V/D409VvsWT_ = .021, Mann-Whitney *U*). Conversely, some of the recorded hypotrophic changes were associated with the plastic changes in the brainstem’s catecholaminergic neurocircuitry (Supplement, **Figure S8b-d**). More specifically, in the dopaminergic region of the ventral tegmental area (VTA) and substantia nigra pars compacta (SNpc), a reduction in tyrosine hydroxylase-positive (TH+) area was recorded (D409V/WT:13.332±2.708%;D409V/D409V:12.538±1.171%;WT:14.666±1.017%; *P*_D409V/D409VvsWT_=.027, Mann-Whitney*U*).

Overall, the observed changes in D409V/WT mice followed the same trend towards the volume loss or gain in all regions of interest (ROIs) as recorded for D409V/D409V (Figure 2B), suggesting a ‘dose-effect’ of the GBA-mutation on putative neurocircuitry phenotypes. However, D409V/WT-specific structural changes were the most prominent in the regions of medial geniculate nuclei, thalamus and RN, with further distinct cellular changes localised to the noradrenergic region of subcoeruleus/sublaterodorsalis nuclei A6sc(*P*_D409V/WTvsWT_=.046, Mann-Whitney *U*; Supplement **Figure S8c**).

In summary, we here demonstrate for the first time a striking RBD-like EEG phenotype^3, 4^ in GBA-mutation (D409V/WT) mouse carriers, accompanied by specific structural changes in the more widespread sleep neurocircuitry (Figure 2). Moreover, our findings are in keeping with human data for patients with GBA-mutation where increased grey matter changes in the brain regions of DRN, midbrain, midbrain reticular formation and PPT have been recently reported^7^.

Of particular note to our study are also findings of two very recent studies^6, 13^. The preliminary results of the first study, conducted in RBD patients, successfully argue for microstates of REM (e.g. phasic *versus* tonic REM) as a so far ignored, and yet likely highly informative and relatively widely accessible, EEG biomarker of RBD^6^. In a second study, Ramirez-Villegas and colleagues (2021)^13^ elegantly argue for brainstem nuclei (e.g. parabrachial and SLD; Supplement **Figure S8**) as pivotal generators of the two types of ponto-geniculate-occipital (PGO) waves that subsequently direct widespread cortical NREM-REM sleep state changes^13^. It is here also suggested that putative mechanisms for such selective widespread neuronal modulations might be twofold, via cholinergic neuromodulation associated with the ascending brainstem–hippocampal synchronizing pathway that terminates in the medial septum and diagonal band of Broca, both shown as affected in our mice (Figure 1), and secondly via direct pontine input to the hippocampus^13^. In light of these two studies, our observation of a reduction in REM-derived theta, with similar qualitative change in NREM-REM band transients in D409V/WT mice (Figure 1), could be taken to reflect underlying changes in two types of PGO waves^13^. Moreover, this possibly also suggests an already defective pontine coordination mechanism in D409/WT mice, with likely aberrant systems and synaptic memory consolidation, as well as synaptic homeostasis. This notion is arguably also supported by our distinct neuroanatomical and histological findings that are further suggestive of an already altered function of cholinergic and noradrenergic brain networks (Figure 1; Supplement).

Despite obvious limitations of our preliminary and cross-sectional study, we believe that the findings posit important translational questions in regards to whether the reported changes in REM-sleep and its theta-frequency-range power^14^, along with involvement of core circuitry structures such are PPT, RN and SLD suggest a differential phenotype of any early non-motor (e.g. cognitive, memory and mood) symptoms in non-manifesting human GBA carriers. Similarly, we believe that these findings also advance the relevance of the GBA1 mice as a valuable model for future exploration of neurostructural and physiological targets in iRBD, and its links with other α-synucleinopathies.

## Methods and Materials

Three mouse lines were investigated, heterozygous (D409V/WT), homozygous (D409V/D409V) GBA1 mutant mice (D409V/WT) and wild type (C57Bl/6), (further in the text D409V/WT, D409V/D409V and WT respectively), all previously described by us and others^12, 15^(**Supplement Methods; Figure S1** depicts the research protocol). D409V/WT (heterozygous GBA1 mice) carry one copy of the human D427V point mutation (human equivalent of the murine D409V point mutation) in the murine *glucocerebrosidase* (*Gba*) gene. Video-electroencephalogram(EEG)-recordings and polysomnography investigations were done according to a strict twelve-hour light-dark cycle protocol, as previously described^14^. All investigations were performed in accordance with the United Kingdom Home Office Animal Procedures Act (1986) and approved by the King’s College London Animal Welfare Ethical Review Body (AWERB).

### Sleep Recording and EEG Analysis

Ten heterozygous D409V/WT and eight WT mice were implanted with screw electrodes (10–50 kΩ) (Supplement, **Figure S2**). The monopolar screw electrodes record the frontal and parietal EEG with ground at occipital bone. In addition, a pair of stainless-steel EMG electrodes was implanted in the dorsal neck muscle. The electrodes were secured with dental cement and standard EEG was recorded for twenty-four hours. EEG/EMG signals were sampled at 200 Hz, amplified 100× and low-pass filtered at 100 Hz using a pre-amplifier (please refer to Supplement Methods for further in depth description). Sleep scoring was performed manually on ten-second epochs using *Sirenia Sleep* software (*Pinnacle-Technology-Inc*.) and signal processing analysis was conducted, as previously described. All statistical tests were performed in “GraphPad Prism” or IBM SPSS Statistics version 25.0. Kolmogorov–Smirnov test was used for normality. Data are represented as the mean ± SEM, unless otherwise stated. Two-way ANOVA (time and treatment factors) and two-tailed unpaired *t-*test were used for the analysis of the sleep and EEG spectrum power data. *P* values are shown when they are less than 0.05.

*(please refer to Supplement for further in-depth description and the full list of
methodological references).*

### Neuroimaging Methodology

#### MRI and Statistical Analyses

The MR images were acquired on 9.4T scanner from *ex vivo* brains and processed using a combination of FSL, ANTs and the QUIT toolbox, as previously described by our group^16^. Tensor based morphometry (TBM) was used to assess the effect of genotype on regional brain volumes. A group comparison was carried out on the Jacobian determinant images with permutation tests and Threshold-Free Cluster Enhancement (TFCE) using FSL randomise. Voxelwise differences data (Supplement, **Figures S4-6**) are displayed on the mouse template image. We also performed a region of interest (ROI) analysis using 71 anatomical ROIs derived from a modified Allen brain atlas, that we co-registered to the study template used for TBM. Volumes of each ROI were calculated by summing the Jacobian determinants within the ROI, and univariate pairwise group comparisons performed using Mann-Whitney U-tests. *(please refer to Supplement for the full list of methodological
references).*

Further detailed methodological description of experimental and study procedures, including the undertaken statistical and MR analyses and all the pertinent methodological references, are available in the **Supplement**.

## Acknowledgments

This work was supported by the Wellcome Trust’s [103952/Z/14/Z] and [212934/Z/18/Z]. KI wishes to thank The Croatian Science Foundation grant: IP-2016-06-8636 („Neuroreact”, PI:SKB).

This work is dedicated to people of Petrinja (Croatia) who suffered through a series of devastating Earthquakes in the harsh winter 2020, in the height of Covid-19 pandemic. Their courage, selflessness, kindness and comradeship are inspiring, long may it continue.

## Conflict of interest statement

C.G., D.C., P.F. and I.R. designed the study. C.G., D.C., C.S., E.K., K.I. conducted and analysed data, all authors were involved in preparation of the manuscript. The authors declare that the research was conducted in the absence of any commercial or financial relationships that could be construed as a potential conflict of interest.

## Data availability

All data that support the findings of this study are available upon reasonable request from the corresponding author.

## Notes

### Competing Interest Statement

The authors have declared no competing interest.

