## Supplement for "Dispersed Sleep Microstates and Associated Structural Changes in GBA1 Mouse: Relevance to Rapid Eye Movement Behavior Disorder"

*joint authors

Running Title: GBA1 mouse and REM sleep

### Supplement Outline

1. Materials and Methods

- Figure S1 – Research Protocol

- Figure S2 – PSG and EEG set up

2. Electroencephalogram and Sleep Changes

- Figures S4

3. Structural Neuroimaging Changes

- Figures S4-S7

4. Histologic Ultrastructural Differences

-Figure S8

*References*

### 1 Methods and Materials

**1.1 Animals:** Three mouse lines were investigated, heterozygous (D409V/WT), homozygous (D409V/D409V) GBA1 mutant mice (D409V/WT) and wild type (C57Bl/6), aged 11.06±0.75 months and further in the text (D409V/WT, D409V/D409V and WT respectively), all previously described by us and others[1, 2] (see Figure S1). D409V/WT (heterozygous GBA1 mice) carry one copy of the human D427V point mutation (human equivalent of the murine D409V point mutation) in the murine *glucocerebrosidase* (*Gba*) gene.

***Experimental Design:*** *All investigations were performed in accordance with the United Kingdom Home Office Animal Procedures Act (1986).*

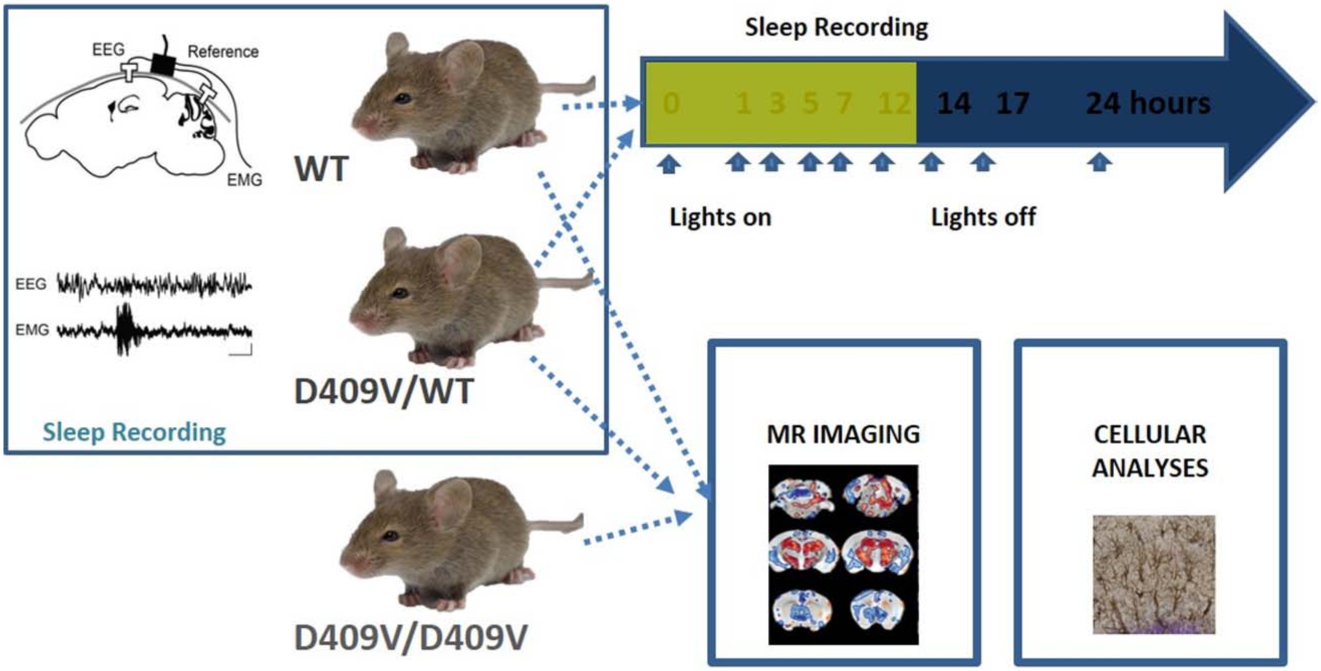


***Figure S1*** **Schematic presentation of the research protocol.** Three mice groups were compared in this study: D409V/WT, D409V/D409V and WT, aged 11.06±0.75 months. EEG sleep recordings were initially performed on age matched D409V/WT and WT mice. Subsequently, imaging and histologic investigations were conducted on all three groups D409V/WT, D409V/D409V and WT. Adapted from[3].

***1.2 Surgery and EEG recording:*** Video-electroencephalogram (EEG) recordings and polysomnography investigations were done according to a strict 12-hour light-dark cycle protocols, as previously described[4].

EEG sleep recordings were performed on ten D409V/WT (eight male) and eight WT (six male) mice. For tethered electromyogram (EMG) and EEG recordings, mice were chronically implanted with skull screw electrodes (Figure S1) to measure cortical EEG. A pair of stainless-steel EMG electrodes was implanted in dorsal neck muscle. Electrodes were connected to head-mounts and secured with dental cement. The animals were allowed two weeks to recover from surgery before the experiments were performed. At the time of the recordings, mice were tethered to four channel EEG/EMG recording systems (*Pinnacle Technology Inc.*) for 30 hours. Recordings took place in a chamber that was programmed with the same light/dark cycles, temperature and humidity equivalent to their home cages. EEG/EMG signals were sampled at 250 Hz, amplified 100× and low-pass filtered at 100 Hz. A two EEG channel and two EMG channel mouse pre-amplifier was used (*Pinnacle Technology Inc*).

***1.3 Sleep Scoring***: Sleep scoring was performed manually on ten-second epochs using *Sirenia Sleep* software (*Pinnacle Technology Inc.*). EEG and EMG recordings were synchronized for each epoch to video recordings. Epochs with EMG amplitude slightly (quiet WAKE) or significantly higher than baseline (active WAKE), together with desynchronized low amplitude EEG were scored as "WAKE". Epochs with low-amplitude EMG and high amplitude delta (1-4 Hz) activity were scored as "NREM" and epochs with low amplitude EMG accompanied by low-amplitude rhythmic theta activity (6-9 Hz) were recorded as "REM" (Van Gelder, 1991). Further analysis of EEG data in the frequency domain was performed using Fourier transforms. Spectrograms showing the amplitude of EEG signals in the time and frequency domain were generated using either Morlet wavelet[5] or using short-time Fourier transform with 1024 samples window length, as previously described[6]. EEG signal analyses were conducted using custom codes written in MATLAB, as previously described[6-8].

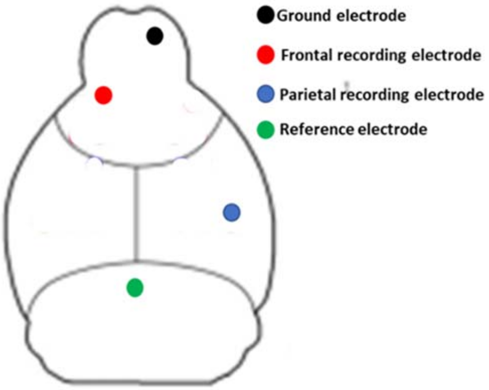


***Figure S2***: **Location of reference, ground and recording electrodes.** Screw electrodes were placed in burr holes in the skull over the parietal cortex (–1.5 mm Bregma, +1.5 mm midline) and frontal cortex (+1.5 mm Bregma, –1.5 mm midline) with a reference electrode over the cerebellum (1.0 mm caudal to lambda, 0 mm midline) and a ground over the olfactory bulb area.

***1.4 Neuroimaging Methodology***

*MRI Acquisition:* Ex vivo MRI was performed on 9.4T Bruker BioSpec 94/20. Perfusion-fixed mouse heads, rehydrated for at least 20 days in phosphate buffered saline (PBS) with 0.05% sodium azide, were placed four at a time in a 50 ml Falcon tube filled with fomblin (*Solvay*) and scanned using a 39-mm birdcage RF transceiver coil. T2-weighted images were acquired using a 3D fast spin-echo sequence: effective echo time

30 ms, repetition time 3000 ms, field of view 25×25×20 mm, acquisition matrix 250×250×200, scan time 5 h 44 m.

*MRI and Statistical Analyses:* The MR images were processed using a combination of FSL[9], ANTs[10] and the QUIT toolbox[11], as previously described by our group[12].

To this end, a study-specific template was created from the T2-weighted images using ANTs (*antsMultivariateTemplateConstruction2.sh*). All subjects were then registered to the template via sequential rigid, affine, and nonlinear (*SyN-algorithm*) registrations (*antsRegistration*). Maps of the Jacobian determinants of the nonlinear deformation fields were computed and log-transformed (*ANTs CreateJacobianDeterminantImage*). The Jacobian determinant is the voxel-wise ratio of volumes between an individual subject and the study template.

Voxel-wise group analysis was then carried out on log-Jacobian determinant images with permutation tests (5000 permutations) and Threshold-Free Cluster Enhancement (TFCE) using FSL randomize[13, 14]. Voxelwise differences data (Supplement, Figures S4-6) are displayed on the mouse template image, using the dual coding approach: differences are mapped to colour hue, and associated P-values are mapped to colour transparency. Contours delineate statistically ($P<0.01$) significant differences. The Allen mouse brain atlas (common coordinate framework version 3), modified in-house to comprise of 71 anatomical ROIs, was similarly registered to the study-specific template for ROI-based analysis. Volumes of each ROI were calculated by summing the Jacobian determinants within the ROI, and univariate pairwise group comparisons performed using Mann-Whitney U-tests.

***1.5 Perfusion and Histology For MRI*:** The mice were killed by transcardiac perfusion with ice-cold heparinized saline followed by 4% buffered paraformaldehyde (PFA). The heads were removed, stored in paraformaldehyde (PFA) for 24 hours and then rehydrated in Phosphate Buffered Saline (PBS) with 0.05% sodium azide at -4ºC for a minimum of 30 days[15] before being MR imaged. After scanning, the brains were extracted from skulls, cryoprotected in 30% sucrose and sectioned at 35µm in a series of six, on a freezing microtome and stored free floating in cryoprotectant at -20°C. For each antibody one series was washed 3 x five min in TBS before incubating in 1% H2O2 for 15 mins, then 10% skimmed milk powder and finally incubation in anti-IBA-1 antibody (1:2,500, Alpha Labs UK (019-19741) Rabbit polyclonal) or anti-tyrosine hydroxylase antibody (Millipore AB152 Rabbit Polyclonal, 1/2000) overnight at 4°C. This was followed by incubation in a biotinylated secondary antibody (anti-rabbit in goat, 1:1000, Vector Labs (BA-1000)) for two hours and ABC kit for 1hr (Vectastain ABC Kit, Vector Labs (PK-6100)). Washes with TBS-X (3 x five min) were performed between each step. Staining was visualised using 3,3′-Diaminobenzidine and sections were mounted onto slides, air dried, dehydrated in increasing concentrations of IMS (70-100%, five minutes each), defatted in Xylene and cover slipped.

The sections were scanned at x40 magnification on a Leica SCN400F slide scanner. Analysis of Iba1 positive, microglial cells was conducted using the optical fractionator method, as previously described [16, 17].

*Analysis of tyrosine hydroxylase (TH) positive area and TH immunoreactivity.* All preselected regions of interest, VTA, SNpc, Raphe nuclei, PB, A6sc, A7, were selected, as previously described[18]. Equal size region was defined on all brain slides with TH positive staining. After adjusting the threshold, TH positive area was measured. Area is measured as % of selected ROI. After calibrating all images (ImageJ software) integrated optical density was measured in all regions of interest, as previously described[19]. All regions (VTA, SNpc, Raphe nuclei, PB, A6sc, A7) were selected using freehand tool.

TH and *iba1* staining (see Figure S8) were analyzed by the thresholding method, as previously described by our group and others[20-22]. Briefly, using an open source *Aperio Image Scope* image viewer, the ROI were set at 4x magnification and a trichromatic image was generated with predefined thresholds for color saturation. The images were subsequently converted to eight-bit *BW* images and analyzed using *ImageJ* (1.51m9; https://imagej.net/ImageJ). The surface area fraction was generated, as percentage of pixels in the image that remained after applying *Image J's* Huang threshold and background subtraction, as previously shown[20, 21].

***1.6 Statistical Analyses***

*Polysomnography*: All statistical tests were performed in "GraphPad Prism". Kolmogorov–Smirnov test is used for normality. Data are represented as the mean ± standard error of mean (SEM), unless otherwise stated. Two-way ANOVA (time and treatment factors) and two-tailed unpaired *t*-test were used for the analysis of the sleep data and power spectrum data. *P* values are shown when they are less than 0.05. *Histology*: IBM SPSS Statistics version 25.0 has been used in all statistical procedures. Kolmogorov-Smirnov test has been used to assess data normality distribution. According to the results and small sample size, appropriate non-parametric test was used in following analysis. Differences between WT, D409V/WT and D409V/ D409V groups were analysed with Kruskal-Wallis test with additional *post-hoc* analyses with Mann-Whitney U test (for difference assessment between each two groups). All P values were exact, two sided, and all values below 0.05 were considered significant.

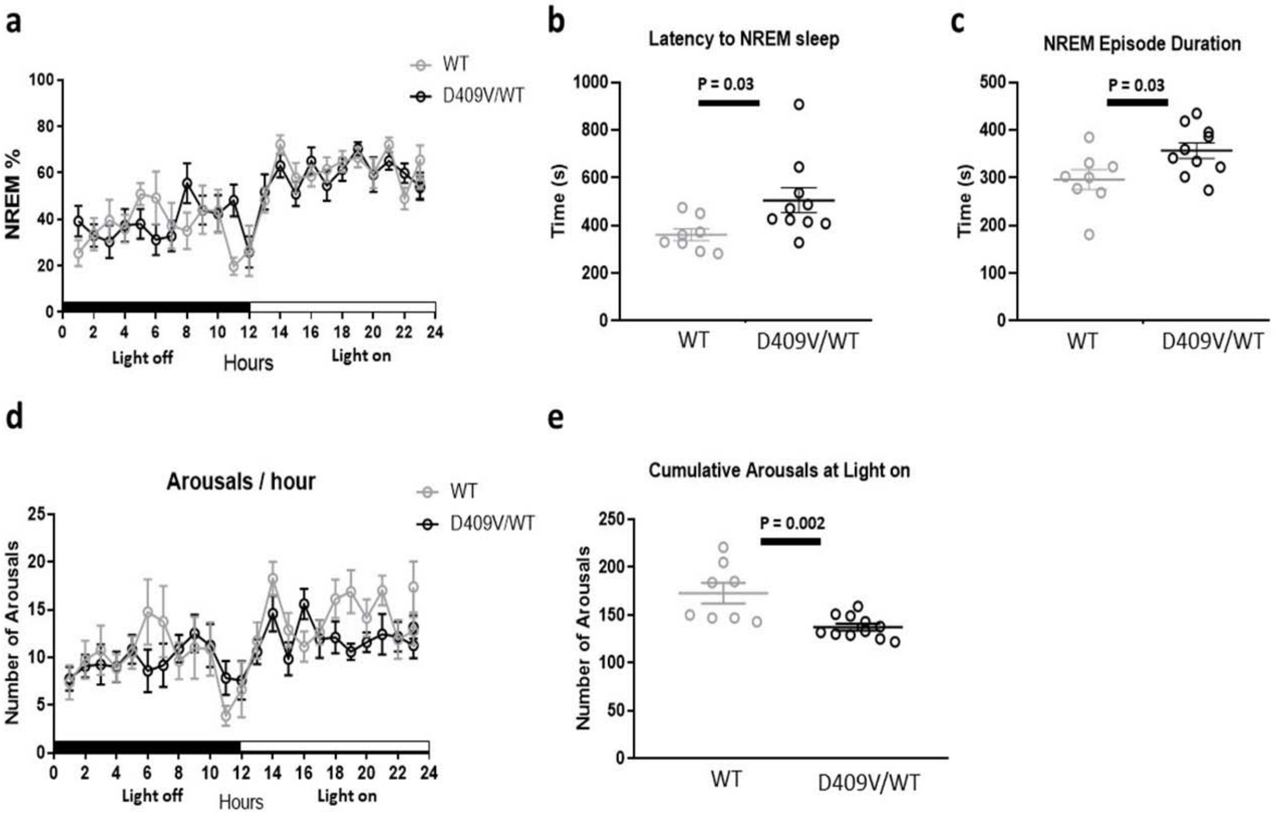


***Figure S3:*** **NREM sleep microarchitecture is affected in mice carrying D409V mutation.** 24 hours scoring of WT and D409V/WT mice showed an overall similar time spent in NREM sleep (a). While time to NREM sleep (sleep latency) was longer (*P*=.03) (b), D409V/WT mice spent longer time in NREM sleep as shown by an increased duration of the average NREM episode (*P*=.03) (c). The increase in the average NREM bout duration was paralleled by a decrease in the number of arousals (*P*=.002) (d,e).

Voxelwise differences showed widespread changes in D409V/D409V versus WT mice, comprising of both increases and decreases (see Figures S4,6,7), whereas the areas of significant changes were more limited in D409V/WT vs WT (Figure S5). To shed further light on these areas of change in an anatomically relevant manner, we next applied an atlas-based analysis (Figure 2) comparing the volume of 71 ROIs between the three genotypes. Comparison of D409V/D409V vs WT mice revealed decreases in the subcortical grey matter of medial geniculate, pretectal, zona incerta and thalamus; also in the white matter tracts of corpus callosum and fimbria fornix, as well as the ventricular volumes (third and lateral).

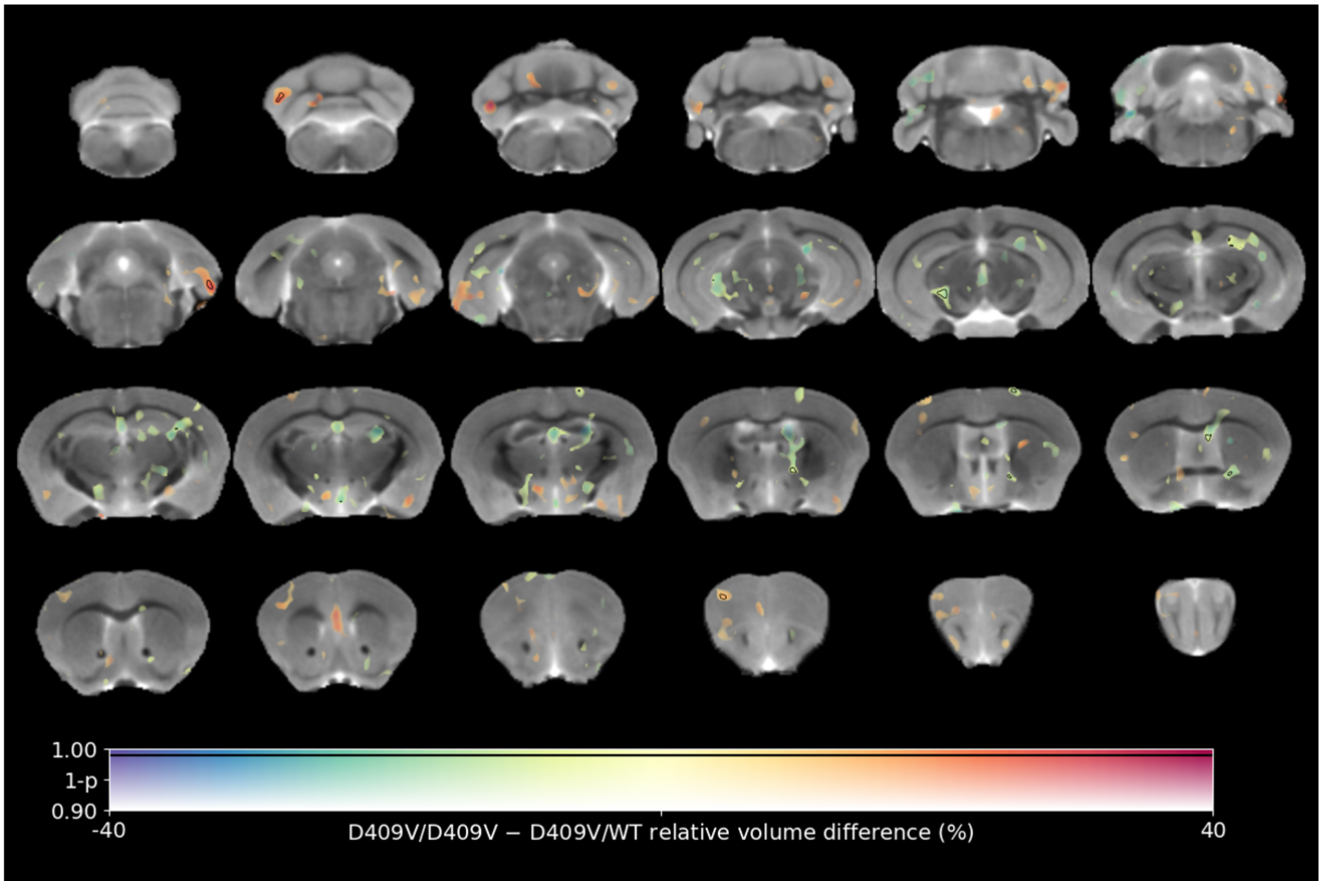


*Figure S4.* **MRI template with an overlay representing relative volume differences (%) in D409V/D409V relative to D409V/WT.** The color of the overlay indicates the inter-group volume difference (warm and cool colours represent gain and loss of volume, respectively), while the transparency indicates the statistical significance, ranging from uncorrected $P$ value 0.1 (transparent) to 0 (opaque). Areas in which uncorrected $P < .01$ are contoured in black.

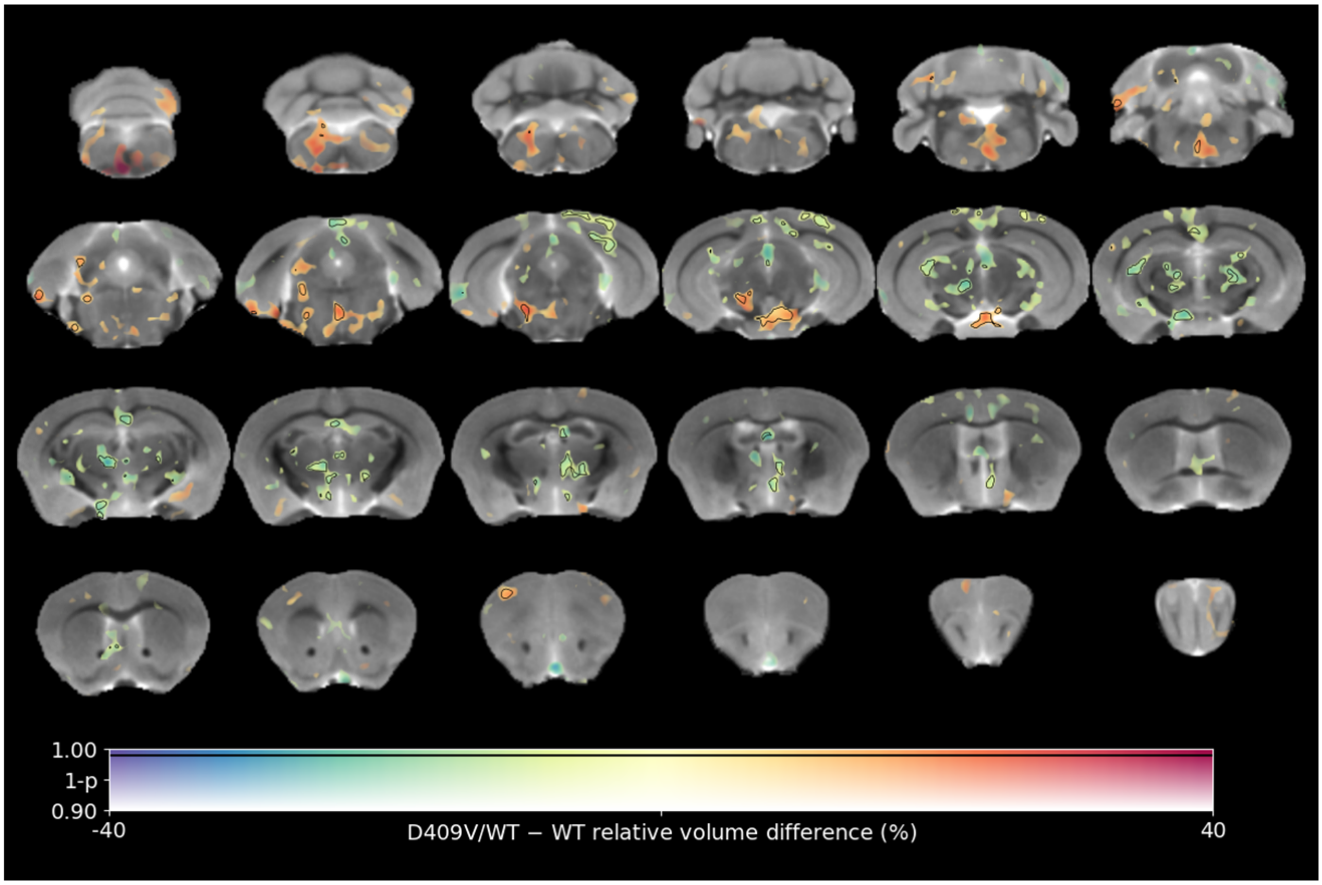


*Figure S5*. **MRI template with an overlay representing relative volume differences (%) in D409V/WT relative to WT mice.** The color of the overlay indicates the inter-group volume difference (warm and cool colours represent gain and loss of volume, respectively), while the transparency indicates the statistical significance, ranging from uncorrected *P* value 0.1 (transparent) to 0 (opaque). Areas in which uncorrected *P* < 0.01 are contoured in black.

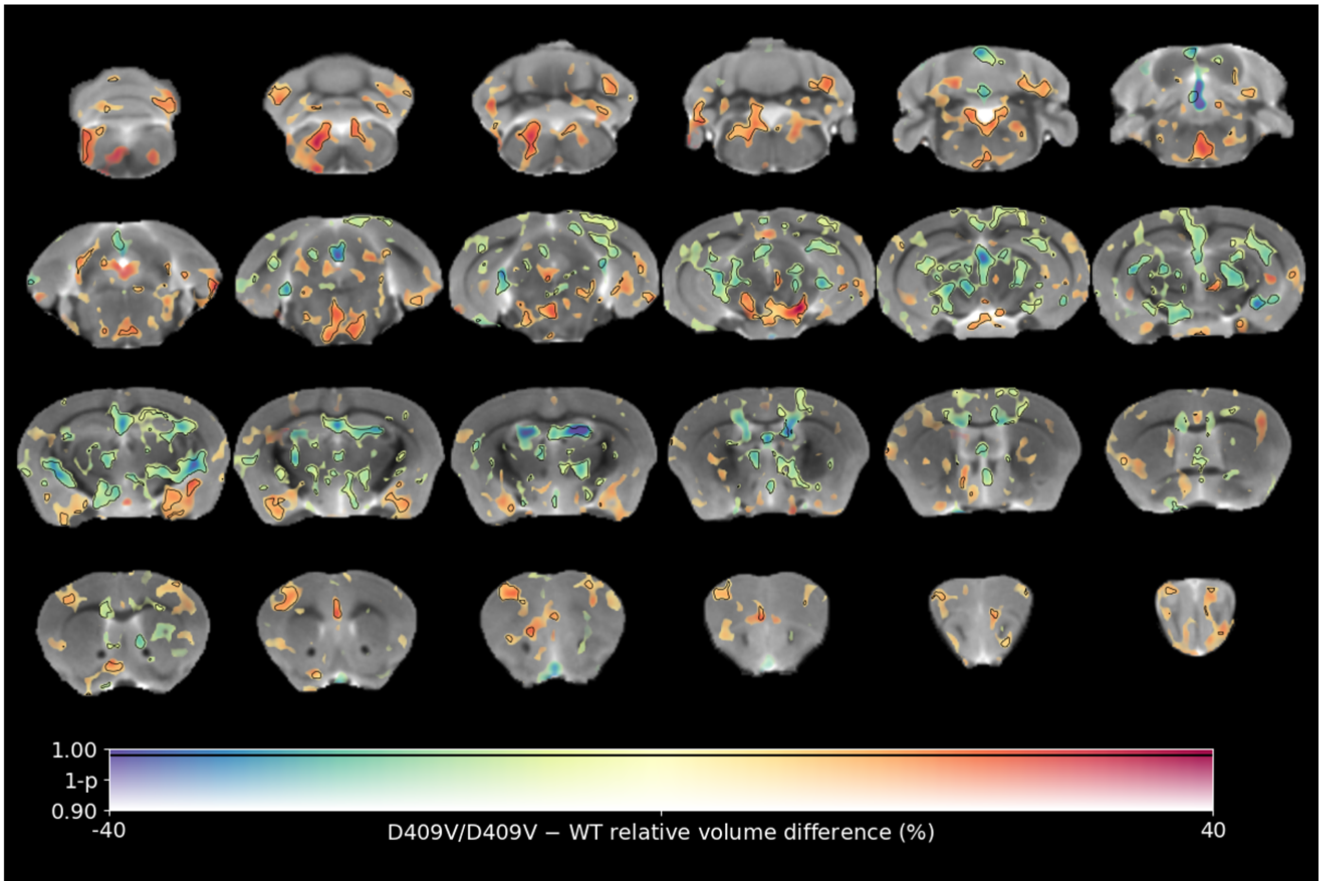


*Figure S6.* **MRI template with an overlay representing voxel-wise volume differences (%) in D409V/D409V relative to WT mice.** The colour of the overlay indicates the inter-group volume difference (warm and cool colours represent gain and loss of volume, respectively), while the transparency indicates the statistical significance, ranging from uncorrected *P* value 0.1 (transparent) to 0 (opaque). Areas in which uncorrected $P < .01$ are contoured in black.

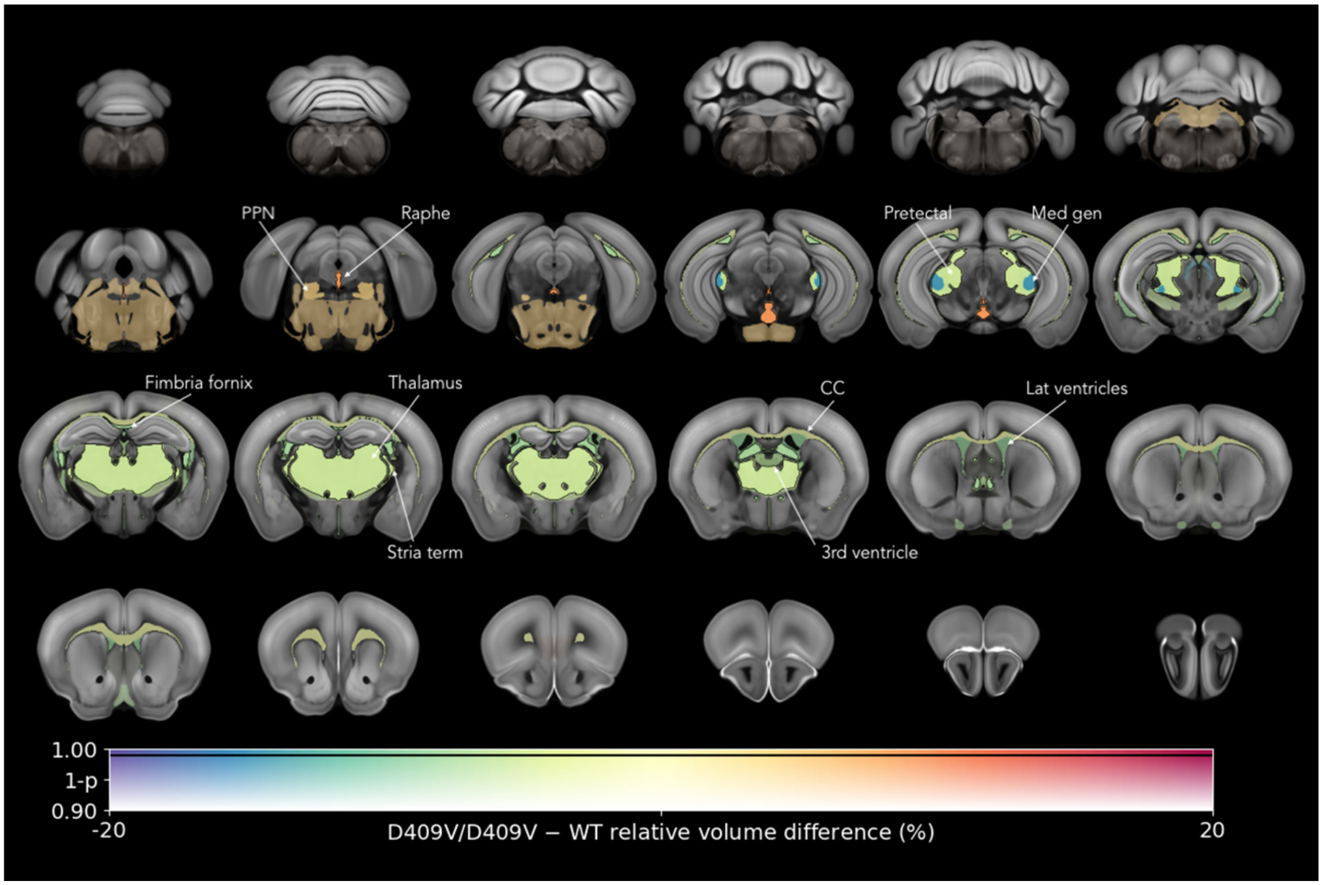


*Figure S7.* **Allen mouse brain atlas template with an overlay representing ROI volume differences (%) in D409V/D409V relative to WT.** The color of the overlay indicates the inter-group volume difference (warm and cool colors represent gain and loss of volume, respectively), while the transparency indicates the statistical significance, ranging from uncorrected P value 0.1 (transparent) to 0 (opaque). Areas in which uncorrected $P < .01$ are contoured in black. *Abbreviations*: PPN – pedunculopontine nucleus; Stria term – stria terminalis, CC – corpus callosum, Med gen – medial geniculate.

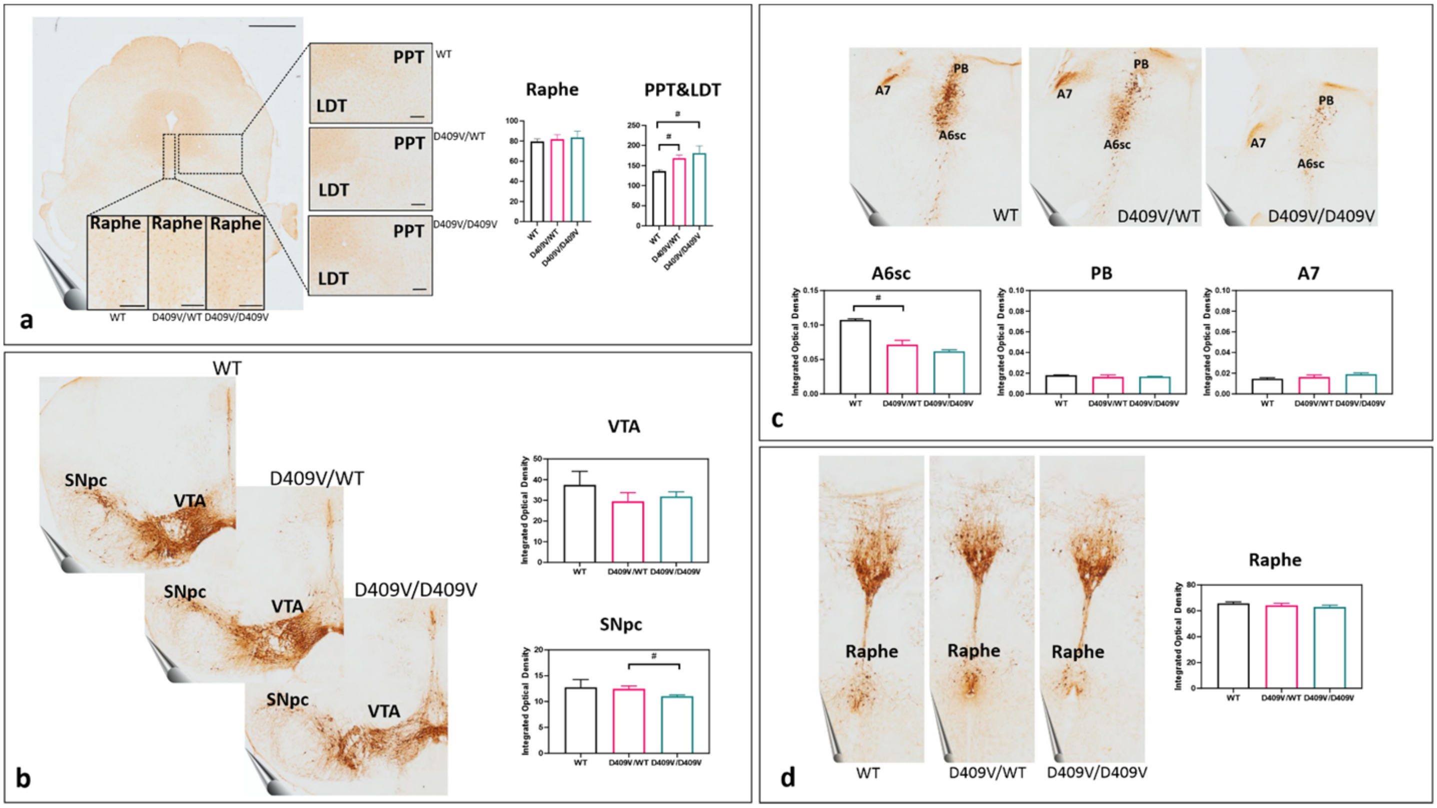


*Figure S8*. **Ultrastructural changes in the regions of interest**. In **a**, representative image of Iba1 staining in PPT&LDT and raphe region is shown. There is an increase in Iba1 positive cells in PPT&LDT region in D409V/WT (*P*= .014) and D409V/D409V (*P*= .021), when compared to WT (Mann-Whitney *U*). There is no significant change in number of Iba1 positive cells in Raphe nuclei region. Scale bar 1 mm, Raphe nuclei 200µm, PPT&LDT 200µm In **b**, representative image of TH staining in VTA and SNpc area is shown. There is no significant reduction in TH immunoreactivity in VTA when D409V/D409V, D409V/WT and WT animals are compared. There is a reduction in TH immunoreactivity in SNpc when D409V/WT and D409V/D409V are compared (*P*= .045, Mann-Whitney *U*). In **c**, a representative image of PB, A6sc and A7 area is shown. There is a decrease in TH immunoreactivity in D409V/WT and D409V/D409V animals compared to WT. The majority of TH signal intensity reduction is observed in A6sc but is only significant between WT and D409V/WT (*P*= .046, Mann-Whitney *U*). Conversely, there is no significant reduction in TH staining intensity in PB or A7. Finally, in **d**, a representative image of TH staining in raphe nuclei is shown. There is no difference TH immunoreactivity signal when WT, D409V/WT and D409V/D409V brains are compared.

*Abbreviations*: VTA – ventral tegmental area, SNpc – substantia nigra pars compacta, PB – medial parabrachial nucleus, A6sc – sublaterodorsal nucleus/locus subcoeruleus, TH – tyrosine hydroxylase, LDT – laterodorsal tegmental nucleus, PPT – pedunculopontine tegmental nucleus, IOD – integrated optical density.

### *References*


1. Clarke, E., *et al.* Age-related neurochemical and behavioural changes in D409V/WT GBA1 mouse: Relevance to lewy body dementia. *Neurochem Int* **129**, 104502 (2019).
2. Sardi, S.P., *et al.* CNS expression of glucocerebrosidase corrects alpha-synuclein pathology and memory in a mouse model of Gaucher-related synucleinopathy. *Proc Natl Acad Sci U S A* **108**, 12101-12106 (2011).
3. Barger, Z., Frye, C.G., Liu, D., Dan, Y. & Bouchard, K.E. Robust, automated sleep scoring by a compact neural network with distributional shift correction. *PLoS One* **14**, e0224642 (2019).
4. Kim, B., *et al.* Differential modulation of global and local neural oscillations in REM sleep by homeostatic sleep regulation. *Proc Natl Acad Sci U S A* **114**, E1727-E1736 (2017).
5. Torrence, C. & Compo, G.P. A Practical Guide to Wavelet Analysis. *Bulletin of the American Meteorological Society* (1997).
6. Zhivomirov, H. On the Development of STFT-analysis and ISTFT-synthesis Routines and their Practical Implementation. . (2019).
7. Gelegen, C., *et al.* Excitatory Pathways from the Lateral Habenula Enable Propofol-Induced Sedation. *Curr Biol* **28**, 580-587 e585 (2018).
8. Gelegen, C., *et al.* Staying awake--a genetic region that hinders alpha2 adrenergic receptor agonist-induced sleep. *Eur J Neurosci* **40**, 2311-2319 (2014).
9. Jenkinson, M., Beckmann, C.F., Behrens, T.E., Woolrich, M.W. & Smith, S.M. Fsl. *Neuroimage* **62**, 782-790 (2012).
10. Avants, B.B., *et al.* A reproducible evaluation of ANTs similarity metric performance in brain image registration. *Neuroimage* **54**, 2033-2044 (2011).
11. Wood, T.C. QUIT: QUantitative Imaging Tools.
12. Wood, T.C., *et al.* Whole-brain ex-vivo quantitative MRI of the cuprizone mouse model. *PeerJ* **4**, e2632 (2016).
13. Smith, S.M. & Nichols, T.E. Threshold-free cluster enhancement: addressing problems of smoothing, threshold dependence and localisation in cluster inference. *Neuroimage* **44**, 83-98 (2009).
14. Winkler, A.M., Ridgway, G.R., Webster, M.A., Smith, S.M. & Nichols, T.E. Permutation inference for the general linear model. *Neuroimage* **92**, 381-397 (2014).
15. Cahill, L.S., *et al.* MRI-detectable changes in mouse brain structure induced by voluntary exercise. *Neuroimage* **113**, 175-183 (2015).
16. Mandarim-de-Lacerda, C.A., Del Sol, M. Tips for Studies with Quantitative Morphology (Morphometry and Stereology). *Int. J. Morphol* **35**, 1482–1494 (2017).
17. West, M.J., Slomianka, L. & Gundersen, H.J. Unbiased stereological estimation of the total number of neurons in thesubdivisions of the rat hippocampus using the optical fractionator. *Anat Rec* **231**, 482-497 (1991).
18. Bucci, D., *et al.* Systematic Morphometry of Catecholamine Nuclei in the Brainstem. *Front Neuroanat* **11**, 98 (2017).
19. Ilic, K., *et al.* Hippocampal expression of cell-adhesion glycoprotein neuroplastin is altered in Alzheimer's disease. *J Cell Mol Med* **23**, 1602-1607 (2019).
20. Westphal, R., *et al.* Characterization of gray matter atrophy following 6-hydroxydopamine lesion of the nigrostriatal system. *Neuroscience* **334**, 166-179 (2016).
21. Abbink, M.R., *et al.* Characterization of astrocytes throughout life in wildtype and APP/PS1 mice after early-life stress exposure. *J Neuroinflammation* **17**, 91 (2020).
22. Khodanovich, M., *et al.* Quantitative Imaging of White and Gray Matter Remyelination in the Cuprizone Demyelination Model Using the Macromolecular Proton Fraction. *Cells* **8** (2019).
